# Investigating tumor-associated macrophages and their polarization in colorectal cancer using Boolean implication networks

**DOI:** 10.1101/2023.08.01.551559

**Authors:** Ekta Dadlani, Tirtharaj Dash, Debashis Sahoo

## Abstract

Tumor-associated Macrophages (or TAMs) are amongst the most common cells that play a significant role in the initiation and progression of colorectal cancer (CRC). [Ghosh et al., 2023] have built a Boolean-logic dependent model to propose a set of gene signatures capable of identifying macrophage polarization states. The signature, called the Signature of Macrophage Reactivity and Tolerance (SMaRT), comprises of 338 human genes (equivalently, 298 mouse genes). The SMaRT signature was constructed using datasets that were not specialized towards any particular disease. To specifically investigate macrophage polarization in CRC, in this paper, we (a) perform a comprehensive analysis of the SMaRT signature on single-cell human and mouse colorectal cancer RNA-seq datasets and (b) adopt transfer learning to construct a “refined” SMaRT signature that specifically characterizes TAM polarization in the CRC tumor microenvironment. Towards validation of our refined gene signature, we use: (a) 5 RNA-seq datasets derived from single-cell human datasets; and (b) 5 large-cohort microarray datasets from humans. Furthermore, we propose the translational potential of our refined gene signature while investigating microsatellite stability and CpG island methylator phenotype (CIMP) in colorectal cancer. Overall, our refined gene signature and its extensive validation provide a path for its adoption in clinical practice in diagnosing colorectal cancer and associated attributes.

**Availability and Implementation:** The data, codes, and software packages used in our research are linked and shared publicly at https://github.com/tirtharajdash/TAMs-CRC.

## 1 Introduction

Colorectal cancer (CRC), characterized by abnormal proliferation of cells within the colon or rectum of the large intestine, is the third most common cancer worldwide, representing 10% of all diagnosed cancer cases [Mármol et al., 2017, Ciardiello et al., 2022]. In 2020, there were an estimated 1.9 million reported cases of newly diagnosed colorectal cancer and 0.9 million deaths globally [Xi and Xu, 2021]. Factors that induce the initiation of the cancer include changes in age, environment, and lifestyle – all of which cause genetic and epigenetic alterations of cells in the gut [Binnewies et al., 2018]. Despite advanced clinical treatments, such as surgical treatment and immunotherapy, CRC is the second leading cause of cancer-related deaths worldwide [Mármol et al., 2017]. As a result, researchers are interested in further exploring the mechanisms and factors that underlie the cancer’s progression and metastasis. The initiation and progression of CRC involve genetic alterations in both the cancer cells and in cells that reside in the surrounding tumor microenvironment (TME). The TME comprises of immune and stromal cells, such as macrophages, dendritic cells, and T lymphocytes — all of which can contribute to chronic inflammation [Binnewies et al., 2018]. Tumor-associated macrophages (TAMs), derived from blood monocytes, are attracted to the tumor site by the activity of growth factors and chemokines within the tumor microenvironment. It is implicated that TAMs are involved in transformation of the normal colonic epithelium to gland-like growths on the large intestine’s membrane, called adenomatous polyps, and to invasive colon carcinoma [Qian and Pollard, 2010].

Macrophages, or MΦ, exhibit distinctive functions in response to the stimuli in their environment; they are traditionally classified as M0 macrophages (unstimulated, undifferentiated), M1 macrophages (reactive-like), and M2 macrophages (tolerant-like). M1 macrophages exhibit pro-inflammatory properties in response to the release of inflammatory cytokines and the presence of oxygen species in the microenvironment. M2 macrophages are characterized by their anti-inflammatory properties, which emerge in response to cytokines produced by immune cells, growth factors released during wound healing, and metabolic changes [Murray et al., 2014, Martinez et al., 2008]. It is conventionally understood that stimuli in a microenvironment can activate M0 macrophages to adopt a tolerant-like or reactive-like polarization state; however, this simplistic nomenclature overlooks the full extent of macrophage plasticity and the continuum of polarization states that macrophages adopt under steady-state conditions and during disease.

[Ghosh et al., 2023] introduced a gene signature, called the Signature of Macrophage Reactivity and Tolerance (SMaRT), which captures the complex dynamics and clinical implications of macrophage phenotype under various contexts. The SMaRT signature was constructed using a pooled human macrophage transcriptome that is represented across different tissues, organs, species, and immune cells. This makes the gene signature a “general” signature that captures the spectrum of macrophage polarization states. In this paper, using publicly available single-cell datasets, we attempt to construct a refined, disease-specific macrophage polarization signature by specializing the SMaRT signature for CRC datasets. Through computational experiments, we demonstrate that our refined CRC-specific signature holds immense potential to be adopted in healthcare research and practice. The overall pipeline of our implementation is provided in Fig. 1.

**Figure 1:**
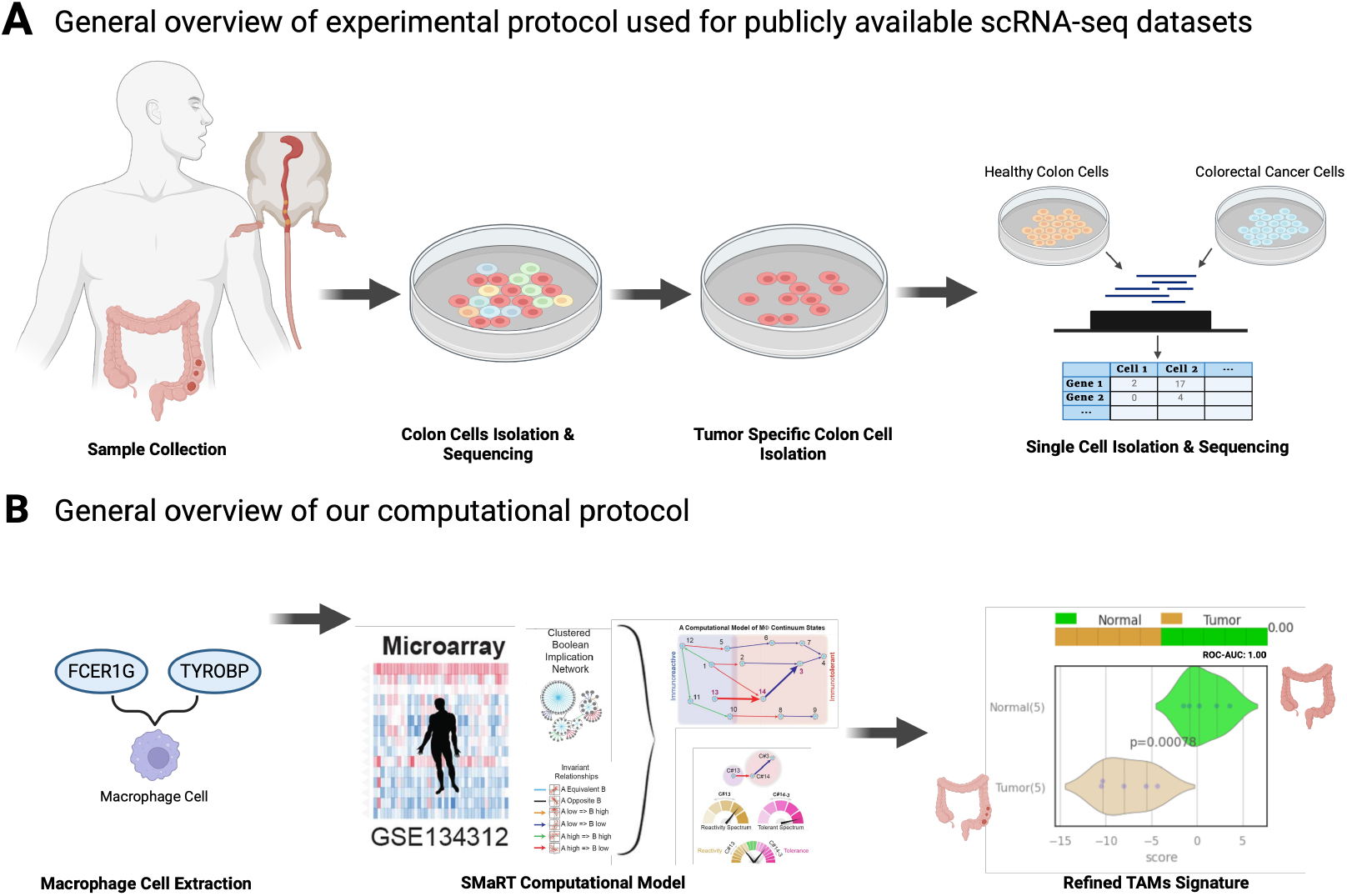
Overview of Experimental and Computational Pipeline: (A) Experimental: To scrutinize the comprehensive cellular dynamics of macrophage polarization in colorectal cancer, we use 8 publicly available human and mice scRNA-seq datasets that sample both healthy colon tissue and colorectal cancer tissue. We utilize an expression profiling matrix provided by high throughput sequencing for each dataset as independent inputs for our computational analysis. (B) Computational: Our computational workflow contains of 3 steps: (1) Extraction of an macrophage cell specific expression profiling matrix from a publicly available scRNA-seq dataset (2) Application of the SMaRT computational model on the macrophage specific matrix and (3) Refinement of the SMaRT signature for a colorectal cancer tumor-associated macrophage specific signature.

Overall, the major contributions of this paper are as follows: (1) We investigate the effectiveness of the original SMaRT signature in capturing the polarization dynamics of macrophage cells, specifically in single-cell RNA-seq colorectal cancer datasets; (2) We propose an idea similar to transfer learning to refine the original SMaRT signature in order to increase the specificity and sensitivity in discriminating colorectal cancer samples from healthy colon samples; and (3) We extensively validate our refined signature to large-scale datasets and demonstrate its translational potential in high-impact clinical studies such as predicting microsatellite stability status and CpG island methylator phenotype (CIMP) in colorectal cancer.

## 2 Results

By applying the SMaRT model to scRNA-seq CRC datasets and refining the generalized SMaRT signature towards the specialized context of colorectal cancer, we investigate and provide computational evidence for the following: (R1) At the single-cell level, macrophage cells in colorectal cancer tissue are characterized by a immuno-reactive polarization phenotype; (R2) Our 24-gene refined SMaRT signature is highly predictive in discriminating between the two sample states: normal, healthy colon (N) and tumorous colon (C); and (R3) Our refined reactive-like gene signature (comprised of 15 genes) has substantial translational potential, primarily validated in determining microsatellite stability status and CpG island methylator phenotype (CIMP).

### 2.1 Differential polarization dynamics of macrophages in colorectal cancer

Fig. 2 shows the polarization dynamics for scRNA-seq dataset GSE132465, which is used for the construction of our refined SMaRT signature. We use statistical significance (*p ≤* 0.05 for a proportions Z-test) when comparing the number of normal and tumorous macrophages in the immuno-reactive quadrant ([C13-low, C14_3-low]) and the immuno-tolerant quadrant ([C13-high, C14_3-high]). We observe a high concentration of tumorous colon macrophage cells in the immuno-reactive quadrant compared to the concentration of tumorous colon macrophage cells in the immuno-tolerant quadrant. This pattern is consistent and statistically significant in multiple scRNA-seq human and mice datasets (see Suuplementary Fig A1). With an increase in tumorigenic activity, TAMs are inclined to convert from the tolerant polarization state to the reactive, anti-inflammatory, cancer-promoting polarization state [Wang et al., 2021]. The reactive polarization state allows TAMs and other tumor cells to promote tumor cell proliferation with the excretion of cytokines; the tumor-promoting phenotype induces the growth and metastasis of CRC cells.

**Figure 2:**
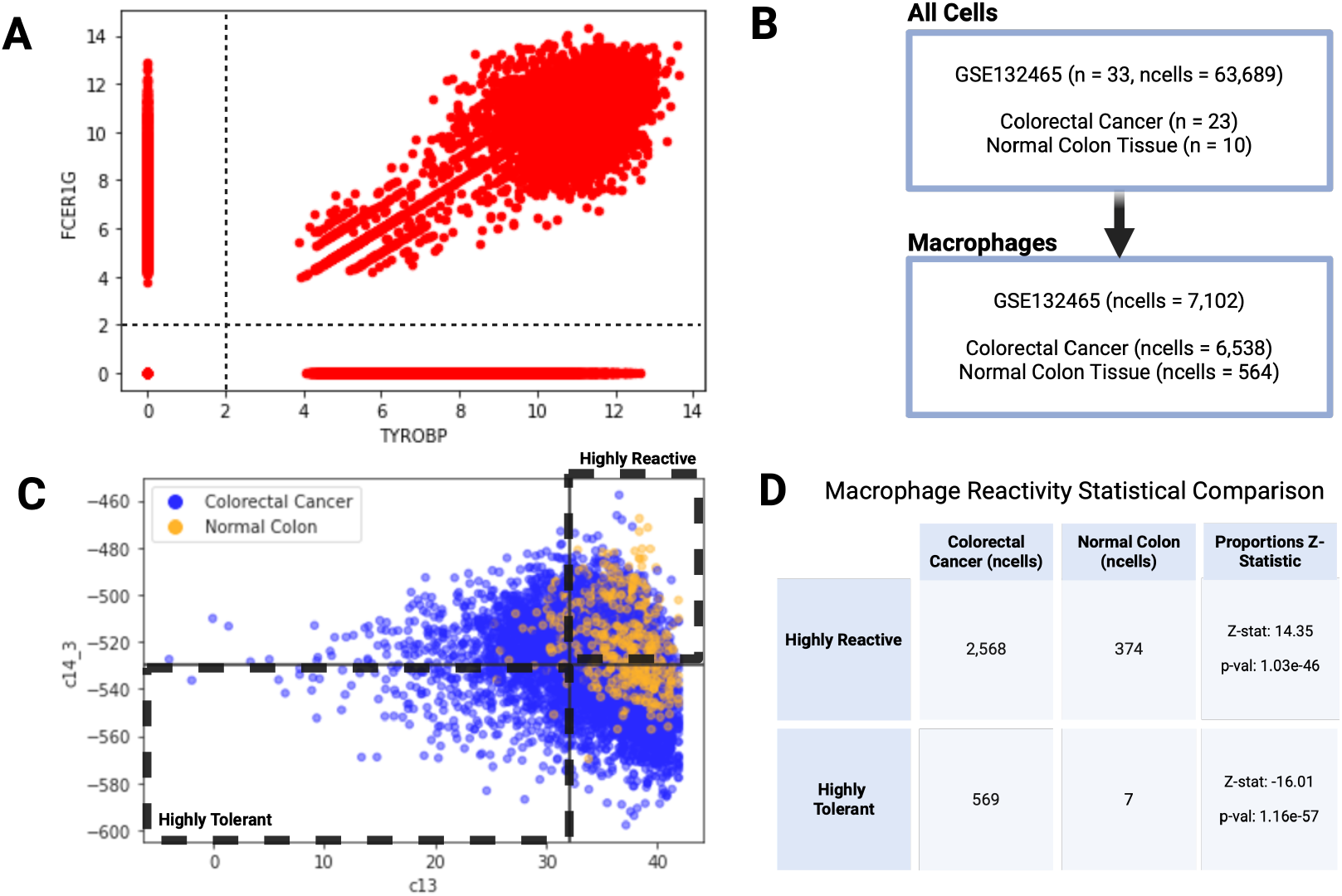
Macrophage polarization dynamics in GSE132465 (scRNA-seq): (A) Scatter-plot showcasing the computational extraction of macrophage cells using thresholds for the expression of universal macrophage biomarkers TYROBP and FCER1G. Each red dot represents a cell from our data. All cells that fall into the upper right quadrant are considered macrophage cells.; (B) Report of the total number of cells used for downstream analysis. About 11% of all cells are considered macrophages.; (C) Scatter-plot comparing the immuno-reactive and immuno-tolerant gene composite scores after application of the SMaRT model to the macrophage-filtered scRNA-seq data. Based on the composite score and the applied StepMiner thresholds, the bottom left quadrant contains highly tolerant macrophage cells and the upper right quadrant contains highly reactive macrophage cells. (D) Statistics comparing the number of highly-reactive and highly-tolerant C and T macrophage cells. There are more highly-reactive colorectal cancer macrophages in comparison to highly-tolerant colorectal cancer macrophages.

### 2.2 Predictive Performance of Refined-SMaRT Signature

Our primary focus in this paper is refinement of the original SMaRT signature. We adopt two different kinds of comparisons while demonstrating the effectiveness of our Refined-SMaRT signature for CRC: (a) a straightforward comparison with the original SMaRT signature, and (b) a comparison with a “noisy” SMaRT signature. Comparison against this noisy signature exemplifies the importance on the necessity for refinement in the context of computational research.

#### 2.2.1 Noisy Signature

We deliberately create a noisy signature with our training dataset by adopting a strategy somewhat opposite to how our refined signature is constructed. 6 *C*13 genes are selected after refinement with filtered macrophages cells separating highly-tolerant N and highly-tolerant C samples and refinement with filtered macrophages cells separating highly-reactive C and highly-tolerant C samples. For the *C*14_3 noisy signature, comprised of 19 genes, we consider all epithelial cells separating N and C samples and all cells separating N and C samples.

#### 2.2.2 Training Dataset Predictive Performance

We now turn to the computational results, as depicted in Figures 3 and 4.

**Figure 3:**
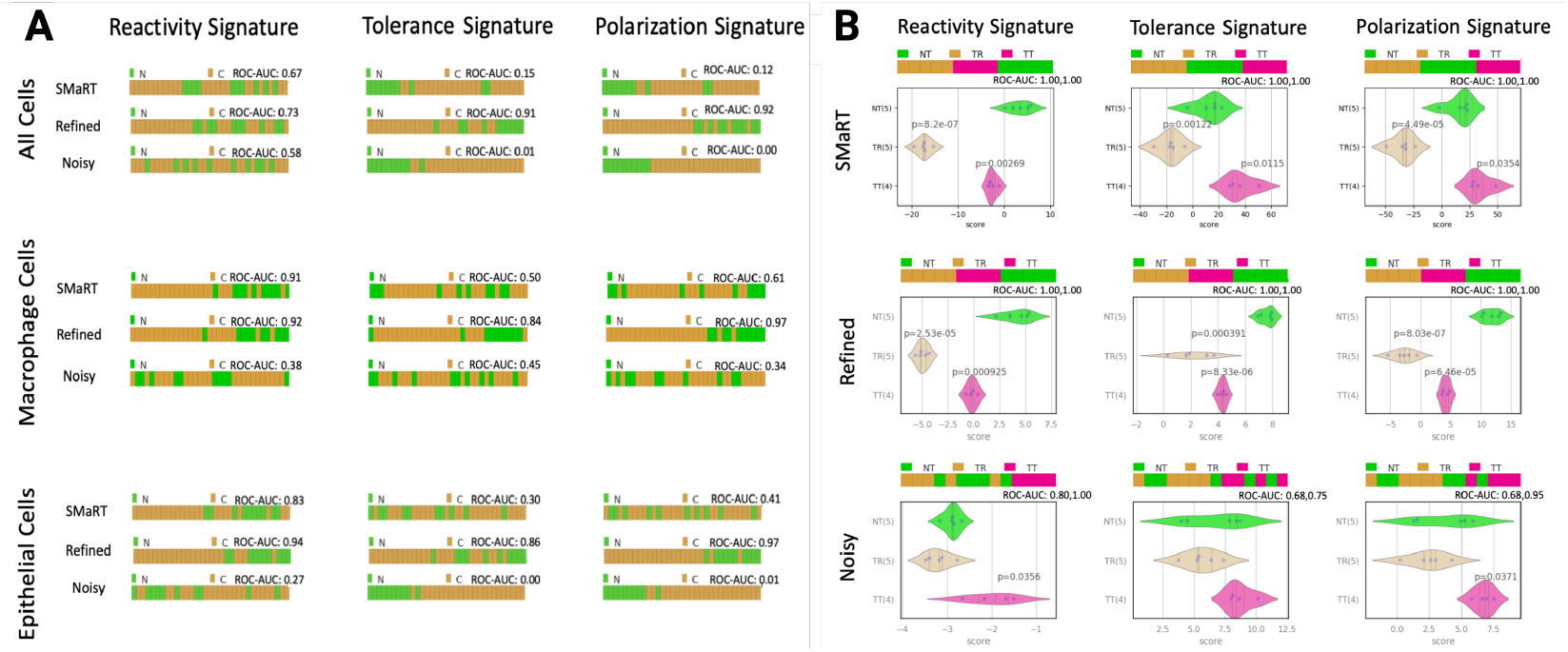
Comparisons of predictive performance between the SMaRT signature, Refined-SMaRT signature, and a noisy signature for the training dataset, GSE132465. Visualizations in (A) report the predictive analysis of the SMaRT, Refined-SMaRT, and noisy signatures for the pseudo-bulk representations in all cells, computationally filtered macrophage cells, and computationally filtered epithelial cells of GSE132465. The violin plots in (B) showcase the predictions of Tolerant NT (NT), Reactive CRC (TR), and Tolerant Tumor (TT) annotations with the SMaRT, Refined-SMaRT, and noisy signatures. The AUC-ROC metric is reported to quantify the predictive potential of the signatures.

**Figure 4:**
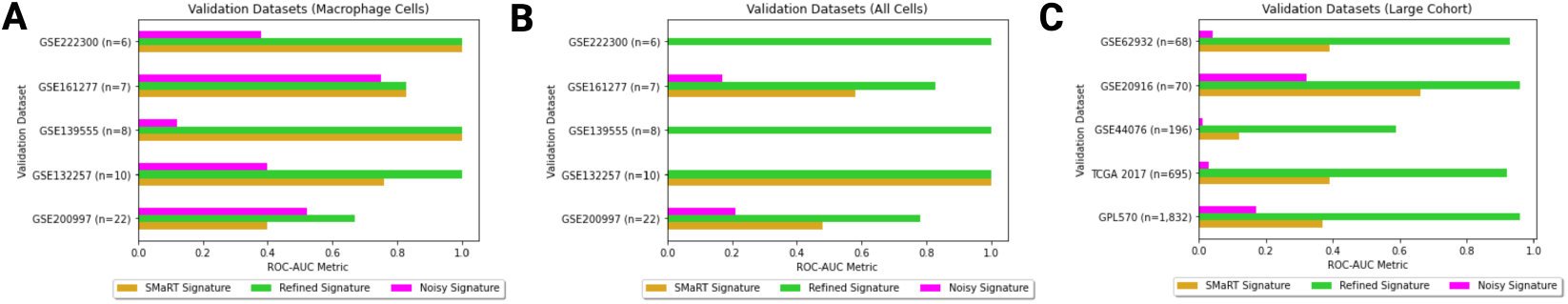
Comparison of predictive performance between the SMaRT signature, Refined-SMaRT signature, and a noisy signature for testing datasets used for validation against pseudo-bulk representations of five human datasets used for scRNA-seq analysis, reported as (A) filtered macrophage cells and (B) all cells. (C) reports validation of the refined signature for 5 publicly-available, large-scaled microarray datasets.

We show the performance of the refined signature on 3 different kinds of pseudo-bulk representations of the training dataset, prepared from all cells, all macrophage cells and all epithelial cells. The results clearly highlight that our minimal 24-gene signature (92% reduction in the number of genes in original SMaRT) has a better predictive capability, with AUC-ROC of more than 90%, in comparison to the original SMaRT signature. Furthermore, the results also suggest that a noisy version of the signature (mimicking any randomly drawn subset of genes from the SMaRT signature) does not have a similar performance.

#### 2.2.3 Validation of Predictive Performance

Validation of the predictive performance of the Refined-SMaRT signature is also reported on the pseudo-bulk representations of the five human datasets used for the preliminary scRNA-seq analysis with the SMaRT signature (See Fig. 4A, B). We compare the predictive performance of the signatures in all cells and in the filtered macrophages. While we report an AUC-ROC range of 0.67–1.00 for the predictive potential, our Refined-SMaRT signature performs better than the SMaRT signature or reports a perfect separation of disease states, similar to the SMaRT signature, while only considering a fraction (7%) of the original gene set. As one would expect, the noisy signature performs worse than our Refined-SMaRT and the original SMaRT signatures, as prominently apparent in the all cells representation of the validation datasets.

Secondary large-scale validation of the predictive performance is showcased in five large-cohort microarray colorectal cancer datasets, in which we do not have the capacity to computationally extract macrophage cells (See Fig. 4C). Analysis in the bulk datasets allows us to validate the signature without the bias of macrophage-specific data. Overall, the current results across datasets are highly indicative of accurate colorectal cancer prognosis and that the robust minimal signature may hold promising implications towards clinical translation.

### 2.3 Reactome Pathway Analysis and Translation Potential

Reactome pathway analysis [Fabregat et al., 2018] is used to identify the enriched signaling and metabolic pathways in our Refined-SMaRT signature. It reveals the potential cellular processes that are impacted by changes in the macrophage polarization dynamics that are likely to occur in the CRC tumor microenvironment as a result of the genes in our signature. We would like to note that many of the genes in our signature are correlated with interferon signaling, which is involved in the modulation of immune responses against pathogens. The activation of reactive macrophages in the tumor microenvironment of CRC tissue, as a response to inflammation, triggers intracellular signaling cascades that up-regulate the interferon-stimulated genes that are involved in immune activation [Min et al., 2022, Chang et al., 2013, Kane et al., 2016].

Microsatellite stability (MSS) and CpG island methylator phenotype (CIMP) are potential markers that differentiate different types of and intensities of colorectal cancer. Microsatellite stability status, studied through the detection of small, repetitive segments of genomic DNA, provides information about how cells handle mishaps in DNA mismatch repairs; its prognostic and immunotherapy predictive status is characterized by two distinct states: MSI and MSS. The stable state (MSS) does not trigger the body’s immune response toward a tumor and typically does not respond to immunotherapy treatments. The unstable state (MSI) corresponds to an unstable tumor and is associated with increased infiltration of immune cells in their microenvironments (i.e., TAMs). Our refined immuno-reactive signature shows that MSI colorectal cancer samples are more immuno-reactive compared to the MSS samples (see Fig. 5B). The higher expression of M1 macrophage genes in the MSI samples suggests that immunotherapy works better with a release of pro-inflammatory cytokines, anti-tumor immunity, and the presence of reactive oxygen species. MSI subtype colon cancer patients have inactivated TGF-*β* and Wnt-*β*-catenin signaling pathways, which are known to reduce the sensitivity to immune therapies (i.e., PD-1 checkpoint blockade therapy) [Wang et al., 2019]. The CpG island methylation phenotype (CIMP) is an epigenetic alteration characterized by the hypermethylation of promoter CpG island sites, resulting in the inactivation or dysregulation of key tumor suppressor genes [Mojarad et al., 2013]. M1 macrophages show a significant effect on the prognosis in CIMP − and CIMP+ (specifically CIMP high) cases [Edin et al., 2012]. Our refined immuno-reactive signature proposes that CIMP+ samples are more immmuno-reactive compared to the CIMP− samples; this supports the relationship between CIMP+ status and MSI status (See Fig. 5C).

**Figure 5:**
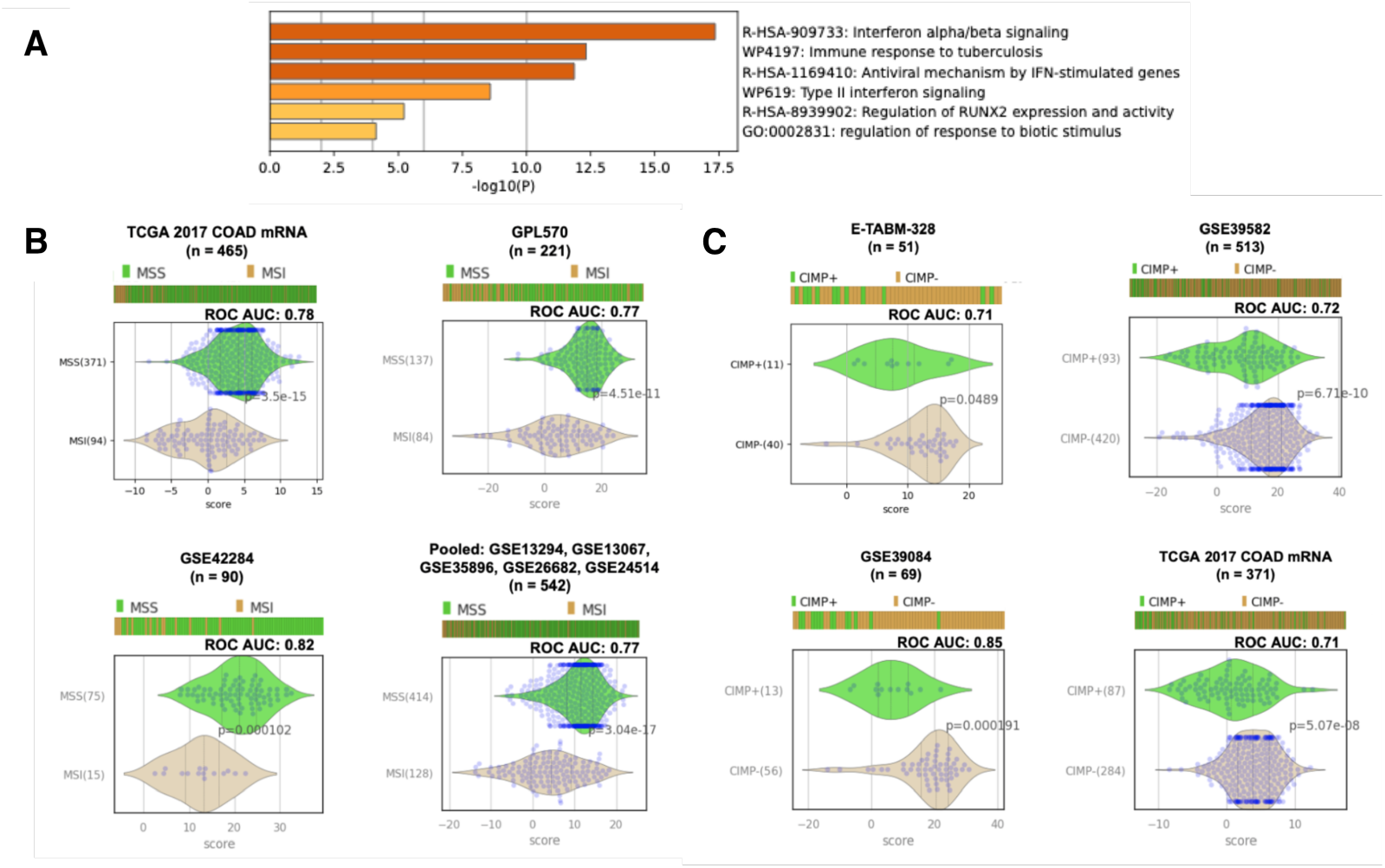
(A) Reactome pathway analysis identifying the biological processes that the genes in our Refined-SMaRT signature are involved in. (B) Predictive potential of our refined immuno-reactive signature towards clinical translation studies in microsatellite stability status. (C) Predictive potential of our refined immuno-reactive signature towards clinical translation studies in CpG island methylator phenotype.

## 3 Discussion

Emerging studies have shown a correlation between the secretion of cytokines and growth factors from tumor associated macrophages (TAMs) that reside in the tumor microenvironment of the colon and the metastatic cascade of colorectal cancer (CRC). TAMs act as promising targets for CRC therapy, precision medicine, and holistic healthcare, because they are often associated with poor prognosis and drug resistance. By adopting a phenotype in the continuum of polarization states between an M1-like and M2-like state, macrophages play a pivotal role in tumor progression and metastasis. Understanding the polarization dynamic and the mechanisms of CRC metastasis would prosper the clinical and prognostic value of TAMs. In this study, we propose a computational technique for the construction of a minimal-and-refined gene signature that captures the polarization dynamics of TAMs in CRC. Our workflow is heavily-dependent on the SMaRT model [Ghosh et al., 2023]—a computational network built to capture macrophage polarization dynamics in a universal manner. We first apply this universal model on scRNA-seq CRC datasets to learn about the general dynamics of polarization states in tumorous and healthy colon macrophages; we find that the TAMs exhibit a immuno-reactive (M1) phenotype. We then develop a refined CRC specific TAMs signature that better captures the polarization dynamics that is reported with the SMaRT-model application on scRNA-seq datasets. Generated based on a t-test dependent differential gene expression method, our 24-gene refined TAMs signature exhibits substantial predictive capability while dealing with a diverse set of CRC datasets and by showing tremendous translation potential in clinical studies. Further extensive examination of our signature is warranted in a wet-lab setup to verify and validate the potential impact for healthcare.

While we found multiple single cell RNA-seq human datasets annotated with colorectal cancer tissue samples and normal colon tissue samples, for which we have a scalable and comparable number of macrophage cells that fell into these two annotations, a lack of publicly available single cell RNA-seq mouse datasets prevented us from developing and validating a mouse CRC TAMs signature. This is because there are not many publicly available datasets with consistent clean boundaries between healthy and cancerous mouse samples, and because within the available datasets, there are a substantially lower number of macrophage specific healthy samples as compared to the cancerous ones. Furthermore, on the computational side, the signature has the potential to undergo further validation with newer CRC-specific datasets, including in spatial transcriptomics data.

## 4 Materials and Method

### 4.1 Data collection and annotation

Eight publicly available single-cell RNA sequencing (scRNA-seq) datasets are collected from the National Center for Biotechnology Information Gene Expression Omnibus (NCBI GEO) database [Barrett et al., 2012, Edgar et al., 2002]. scRNA-seq captures the heterogeneity of RNA transcripts across individual cells; it allows for the understanding of a disease with higher resolution compared to RNA-seq datasets. Gene expression counts, captured in an expression profiling matrix after high throughput sequencing, are normalized by log_2_ scaling of CPM (counts per million) values [Li and Dewey, 2011, Pachter, 2011]. Each sample in a dataset is annotated with metadata, including whether it originates from tumorous colon tissue or normal, healthy colon tissue. A summary of all datasets can be found in Table 1.

**Table 1:** A summary of the publicly available single-cell and microarray datasets used for our study. Usage annotations are as follows: Sc: single-cell analysis; Tr: training towards refinement; Va: validation of refinement; Ta: translational analysis. Superscripts: ^∗^Artificially generated technical replicates; ^*†*^Samples specific to colorectal cancer; ^*‡*^Colorectal cancer specific samples annotated with status for specific translational analysis; ^∗∗^Source datasets: GSE13294, GSE13067, GSE35896, GSE26682, GSE24514

| GEO ID | Species | Type | n-samples | n-cells | Usage |
| --- | --- | --- | --- | --- | --- |
| GSE132465 | *H. sapiens* | Single-cell | 33 | 63,689 | Sc, Tr |
| GSE161277 | *H. sapiens* | Single-cell | 13 | 54,782 | Sc, Va |
| GSE200997 | *H. sapiens* | Single-cell | 23 | 49,859 | Sc, Va |
| GSE132257 | *H. sapiens* | Single-cell | 10 | 18,409 | Sc, Va |
| GSE139555 | *H. sapiens* | Single-cell | 8* | 12,495 | Sc, Va |
| GSE222300 | *H. sapiens* | Single-cell | 6* | 14,852 | Sc, Va |
| GSE198758 | *M. musculus* | Single-cell | 4 | 20,849 | Sc |
| GSE224679 | *M. musculus* | Single-cell | 4 | 27,656 | Sc |
| GPL570 | *H. sapiens* | Microarray | 1832<sup>†</sup>, 221<sup>†</sup> | – | Va, Ta |
| TCGA 2017 | *H. sapiens* | Microarray | 695<sup>†</sup>, 465<sup>†</sup>, 371<sup>†</sup> | – | Va, Ta |
| GSE20916 | *H. sapiens* | Microarray | 70<sup>†</sup> | – | Ta |
| GSE44076 | *H. sapiens* | Microarray | 19<sup>†</sup> | – | Ta |
| GSE62932 | *H. sapiens* | Microarray | 68 | – | Va |
| GSE42284 | *H. sapiens* | Microarray | 90<sup>‡</sup> | – | Ta |
| E-TABM-328 | *H. sapiens* | Microarray | 51<sup>‡</sup> | – | Ta |
| GSE39084 | *H. sapiens* | Microarray | 69<sup>‡</sup> | – | Ta |
| Pooled** | *H. sapiens* | Microarray | 542<sup>‡</sup> | – | Ta |
| GSE39582 | *H. sapiens* | Microarray | 513<sup>‡</sup> | – | Ta |

### 4.2 SMaRT: A machine learning based macrophage polarization model

The SMaRT model [Ghosh et al., 2023] is a computational approach that utilizes the concept of Boolean Implication Network (BoNE [Sahoo et al., 2008]) to segregate the polarization states in macrophages within the immunologic spectrum (M1-and-M2-like phenotype) based on the expression of genes. Below, we first describe the fundamentals of BoNE and then provide information on how SMaRT adopts a good-old-fashioned-AI to construct a machine learning signature to capture macrophage polarization.

Let 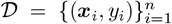 be a labeled dataset of *n* data instances, where each ***x***_*i*_ = [*x*_*i*_, …, *x*_*d*_] represents a *d*-dimensional vector representation (in our case, a vector of gene expression for sample *i*) and *t*_*i*_ is the class-label (in our case, macrophage phenotype: M0-like, M1-like or M2-like). The SMaRT model aims to learn a mapping function: *f*_(*π*,***θ***)_ : ***x***_*i*_ ⟼ *y*_*i*_ such that each gene expression data sample is correctly labeled (with some probability) by *f* as being *y*_*i*_. The function *f* has a structure *π* and some parameters ***θ***. SMaRT adopts the technique of Boolean implication, developed in [Sahoo et al., 2008] and construct a graph-like structure of *f*. From the available training data of gene expressions and macrophage pheno-types, it first derives a robust relational structure to encode the relationships among clusters of Boolean-equivalent genes (genes that are strongly correlated with each other). This relational structure is called *Boolean Implication Network* or BoNE. Each node in the network represents a Boolean equivalent cluster, which consists of a set of genes that have an “equivalent” relationship with each other (strong correlation). An edge between two clusters represents the overwhelming Boolean Implication Relationship (BIR) between the genes in the two clusters. A edge can be labelled of: *low* → *low, low → high, high → low, high → high* and *opposite*, referring to the type of Boolean relationship betwen the two clusters. These relations are derived based on statistical learning on gene expressions within each cluster. We refer the reader to [Sahoo et al., 2008, 2010] for a detailed understanding of the fundamentals of Boolean implications and BoNE. Once a Boolean network has been derived, SMaRT then constructs a parameter called “*composite signature*”, a weighted score to summarize the cumulative expression of all the genes in a BoNE cluster. Mathematically, for example, for a path *C*13 → *C*14 → *C*3 the composite score can be written as:

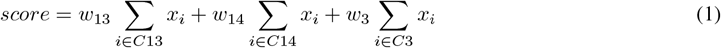

where, *C*13, *C*14 and *C*3 refer to the three clusters, *x*_*i*_ refers to the gene expression value for a gene *g*_*i*_ in a cluster, and *w*_*i*_ refers to the weight paramer for cluster *Ci*. In essence, the SMaRT model involves the following procedural steps to capture the dynamics of macrophage polarization:

#### Step 1. Search

Traverse the Boolean implication network to obtain a path consisting of three nodes with a transitioning relationship: high-low edge (an overwhelming high-low Boolean implication relationship). One way to do is to enumerate out all possible paths with Boolean state transitions and then pick the one that best captures the training data (see below).

#### Step 2. Scoring using a cost-heuristic

Learn the parameters (or coefficients) for the composite signature obtained in Step (1) such that the error in classifying the samples into their corresponding macrophage-polarization state is minimized.

We note the following with regard to the above steps:

1. The idea of clustering equivalent genes results in fewer nodes in BoNE, which allows us to enumerate all possible 3-length paths in the network. This step drastically compresses a network of all possible genes (graph of size *𝒪* (*n*^2^)) to a clustered-gene graph (*𝒪* (*k*^2^)). In real-world scenario, *k* ≪ *n*.
2. Since number of paths is limited with the length of each path (max 3), a simple search over a small range of values (e.g., [− 2, − 1, 0, 1, 2]) for the coefficients (*w*_*i*_s) of the composite signature in Eq. 1 in suffices, without requiring a full optimization-based search which is often resource-intensive.
3. As the cost-heuristic, we use the Area Under the Curve - Receiver Operating Characteristic (AUC-ROC) of the predictive performance to score each path enumerated in Step (1).
4. The final learned model is then the transition path that results in the maximum AUC-ROC, along with the corresponding set of coefficient parameters for the composite score for each cluster.

In the procedure described above, Boolean paths that intersect transitive implication relationships [Sahoo et al., 2008, 2010] between large gene clusters in the BoNE are used to capture the continuum of polarization from the reactive-like pole to the tolerant-like pole of the network. Specific enriched gene paths are selected for testing the classification of samples into the immuno-reactive and immuno-tolerant states in bulk RNA-seq and microarray macrophage datasets annotated with polarization states. Multivariate analysis of the top Boolean paths shows that the path that connects gene clusters *C*13 → *C*14→ *C*3 discriminates the M1-like and M2-like polarization states. Independently, the expression of the genes in C13 accurately predicts the reactivity (M1-like) state and the expression of genes in path *C*14 → *C*3 demonstrates prediction of the tolerant (M2-like) state. The *C*13 → *C*14 → *C*3, or SMaRT, signature consists of 48 human genes (equivalent to 71 mouse genes) in C13 and 290 human genes in the path *C*14 → *C*3 (equivalent to 227 mouse genes). It successfully identifies the M1-like/M2-like polarization states under a diverse range of tissue-resident macrophages, in both human and mice, and in other immune cells.

### 4.3 Single-cell analysis using SMaRT

#### 4.3.1 Macrophage polarization in scRNA-seq datasets

Due to the lack of available macrophage-specific CRC sequencing studies, we apply the SMaRT model on computationally extracted macrophage cells. Filtering is based on the expression of the universal macrophage biomarkers: FCER1G and TYROBP [Dang et al., 2020]. Any expression of FCER1G and TYROBP suggests that a cell is a macrophage; we use previously validated thresholds for the expression of these genes to increase our confidence in filtering TAMs: *FCER*1*G >* 2.0 and *TY ROBP >* 2.0 or *FCER*1*G >* 0.0 and *TY ROBP >* 0.0. The 338-gene SMaRT model is then applied to each filtered scRNA-seq datasets. We plot the gene composite score for the immuno-reactive cluster (*C*13) against the gene composite score for the immuno-tolerant clusters (*C*14 and *C*3) to visualize the polarization dynamics of each macrophage cell in our dataset.

The StepMiner algorithm [Sahoo et al., 2007] is used to binarize these two composite scores into *high* and *low* values. This approach partitions the plotting space into 4 quadrants. Cells that are confined to the [C13-low, C14_3-low] quadrant are classified as *highly reactive* macrophages; and the cells that are confined to the [C13-high, C14_3-high] are classified as *highly tolerant* macrophages. Comparing the number of tumorous and normal macrophage cells that are highly reactive and highly tolerant allowed us to derive a general assumption for colorectal cancer macrophage polarization dynamics. To determine whether the number of tumor-specific macrophage cells and the number of normal colon-specific macrophage cells in a quadrant are significantly comparable, a test for proportions based on a normal (*z*) test is performed. The *p*-value is used to indicate the statistical significance of the two proportions using the *z*-test. That is, *p ≤* 0.05 indicates that we can reject the null hypothesis, which states that there is no difference in the proportion of N samples to C samples.

### 4.4 Construction of a CRC-specific signature using refinement

In this section, we describe a methodology used to refine the “general” SMaRT signature [Ghosh et al., 2023] towards the specialized context of colorectal cancer. The goal of refinement is to remove genes from the SMaRT signature that prevent us from accurately capturing the polarization dynamics of macrophages in colorectal cancer tissue. Mathematically, our refined TAMs signature is obtained from a subset of genes (of the 338 genes in the original SMaRT model) that not only captures the macrophage polarization dynamics but also is predictive towards classifying N and C phenotype. We use the term “refinement” to refer to the steps used in constructing such a signature, which are as follows: We first identify a scRNA-seq dataset where the composite score versus sample disease annotation relationship shows statistically significant outcomes: high-reactive and highly-tolerant quadrants separate the two categories of samples (N and C) very well. In order to derive a refined signature, we manipulate the training dataset into different pseudo-bulk representations based on the provided gene expression matrix and on the conclusions derived from the scRNA-seq analysis pipeline. Here, we would like to note a potential reasoning, owing primarily to domain-knowledge, behind the categorization of cells, we follow subsequently. Due to the capacity and diversity of differentiated cells in the colon [Parikh et al., 2019], we face the issue of potential contamination of “noisy” genes from the SMaRT signature that stem from the colonic epithelium. To ensure that our signature primarily captures macrophage polarization and is not misrepresented by the unique expression pattern of a gene from a differentiated cell, we want to remove the bias of genes that may be expressed differently in epithelial and macrophage cells. Therefore, we consider genes that have similar and consistent expression patterns in both cell types. To computationally extract epithelial cells, a threshold of ≤ 0 for *TY ROBP* and *FCER*1*G* (to remove potential macrophages) and a threshold of *>* 2 for the epithelial biomarker *EPCAM*, which is involved in making the epithelial cellular adhesion molecule (EpCAM) [Goossens-Beumer et al., 2014], is applied.

Since the gene expression levels are measured by the number of reads that map onto each gene, we can perform differential gene expression (DGE) analysis [Liang and Pardee, 2003] by using a *t*-test ranking of the SMaRT signature genes. This approach allows us to compare the statistical significance between our two disease-state annotations (N and C). We expect to capture a pattern that is representative of the results from our scRNA-seq analysis. Specifically, our DGE analysis is performed on 4 types of pseudo-bulk representations (created using log_2_-normalized CPM values) of our training dataset, consisting of:

**Type 1** All cells separating the annotations of N and C samples;

**Type 2** A comparable subset of macrophages cells separating the annotations of highly-tolerant T and highly-tolerant C samples;

**Type 3** A comparable subset of macrophages cells separating the annotations of highly-reactive C and highly-tolerant C samples;

**Type 4** Epithelial extracted cells separating the annotations of N and C samples.

Overlaying the top rankings from the following differential results are used to compose our refined signature:

#### Refining *C*13

Let *n* be the number of genes in *C*13. We take the top *k* genes of *n* expressed in Type 4. Then, of the top *k*, we take the top *m* genes expressed in Type 2.

#### Refining *C*14*_*3

Let *n* be the total number of genes in *C*14 and *C*3 together. We take the top *k* (of *n*) genes expressed in Type 4. Then, of these *k* genes, we take the top *k*_1_ genes expressed in Type 2. Furthermore, we take the top *k*_2_ (of *k*_1_) genes expressed in Type 3. Of the top *k*_2_ genes, we take the top *m* genes expressed in Type 1.

From our scRNA-seq experiments, we choose GSE132465 (*H. sapiens*, n = 33, n-cells = 63,689, n-macrophage cells = 8,049) to be our training dataset. Using the refinement methodology, as described above, we retain 15 immuno-reactive genes (out of the 48 genes in *C*13) and 9 immuno-tolerant genes (out of the 290 genes in *C*14_3). For the refinement of *C*13, we set *n* = 48, *k* = 25, and *m* = 15. For the refinement of *C*14_3, we set *n* = 290, *k* = 116, *k*_1_ = 63, *k*_2_ = 43, and *m* = 9. In summary, our refined signature consists of only 24 genes (in contrast to the 338 genes in the original SMaRT signature):

- Immuno-reactive genes (Refined *C*13): {*OAS*2, *TAP*1, *XAF*1, *IFIT*3, *IFIT*2, *OAS*1, *ISG*15, *TRIM* 21, *MX*1, *APOL*1, *OAS*3, *CXCL*9, *SP*110, *PML, STAT*1};
- Immuno-tolerant genes (Refined *C*14_3): {*RPL*3, *SLC*46*A*3, *CD*302, *RBM*4, *METTL*7*A, CCDC*88*A, RPL*17, *CLEC*10*A, DNASE*1*L*3}.

We use the term “Refined-SMaRT model” to refer to the SMaRT model used to analyze these 24 genes. We calculate a total composite score [Ghosh et al., 2023] to map the expression values of these 24 genes to a single real-value by using a weight of − 1 for the genes in *C*13 and weights of 1 and 2 for the genes in *C*14 and *C*3, respectively. This procedure is similar to building a logistic regression model using the refined set of genes to classify the two classes of samples. We note an interesting observation here: Our process of signature refinement resembles with the way transfer learning [Bengio, 2012, Krishnan et al., 2020] works, that is, to transfer knowledge from a larger (more general) model to a small (more specific) machine learning model.

#### 4.4.1 Validation of the Refined-SMaRT model

The goal of the validation stage is to investigate whether the 24 gene signature obtained in the previous section has any diagnostic benefits. For this, we use two statistical metrics: (1) area under the receiver-operating characteristic curve (AUC-ROC): [Hand and Till, 2001] used to quantify the predictive potential of the composite signature for classifying the N and C samples at the (pseudo)-bulk RNA-seq and microarray levels; and (2) *p*-value: used to determine whether the composite score for the refined gene signature is statistically distinctive between N and C samples. We note that for the scRNA-seq datasets, we create two kinds of pseudo-bulk RNA-seq representations for validation: (a) using all of the cells, and (b) using only the filtered macrophage cells.

## Code and Data Availability

The data, codes, and software packages used in our research are linked and shared publicly at https://github.com/tirtharajdash/TAMs-CRC.

## Acknowledgements

The authors sincerely thank the members of the Boolean Lab for their constructive feedback during this research. This work is supported by the NIH grant R01-AI55696. Other sources of support include R01-GM138385, UG3TR003355, Padres Pedal the Cause/RADY #PTC2017 and #PTC2021.

## Author contributions statement

E.D. and T.D. conceived and conducted the experiments. All the authors analyzed the results. D.S. acquired the funding and supervised the work. All the authors wrote and reviewed the manuscript.

## Appendix

**Figure A1:**
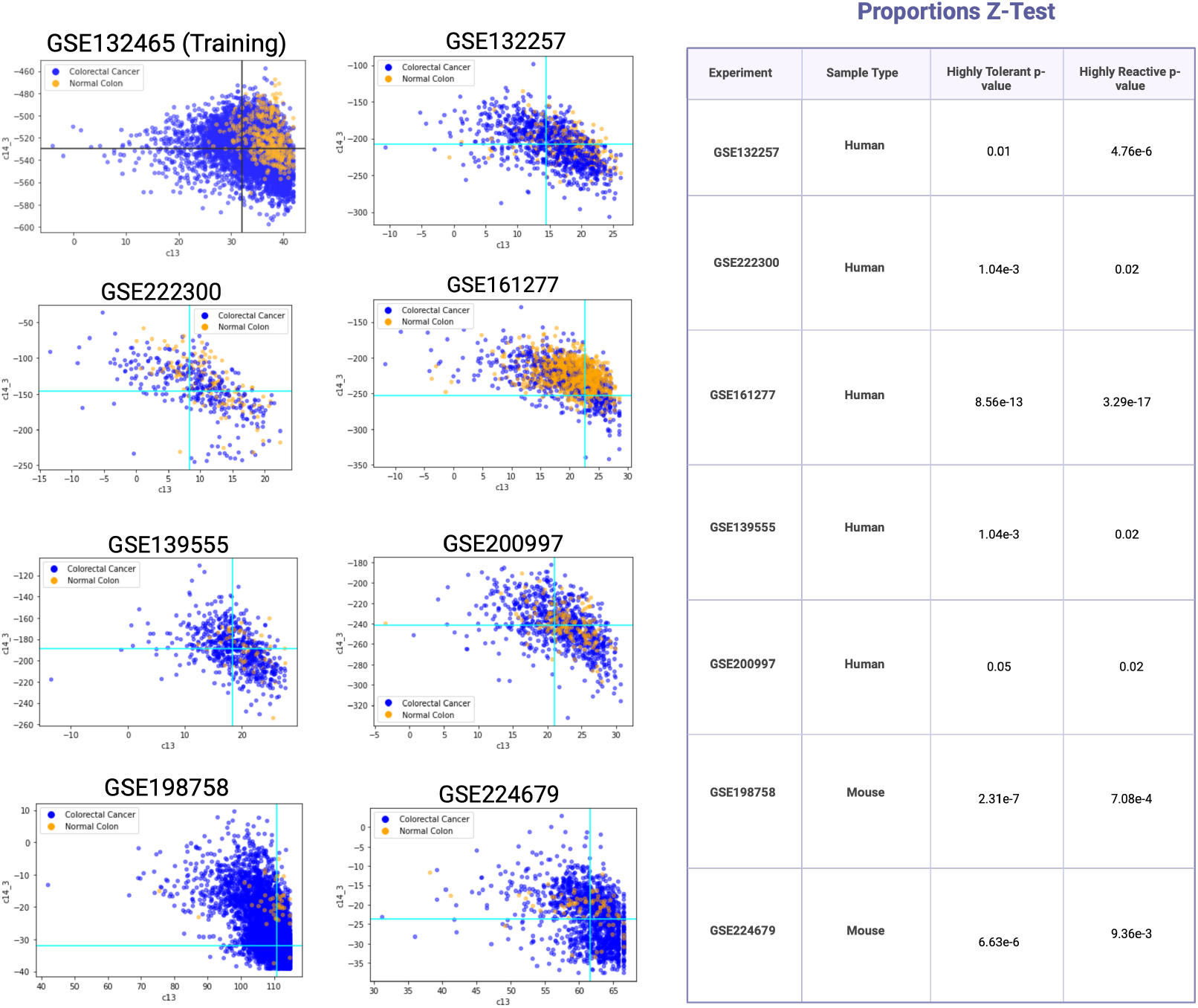
SMaRT Model Application on scRNA-seq human and mice datasets with disease-state annotations (N and C). We see a consistent pattern that there are more highly-reactive C macrophage cells in comparison to highly-tolerant C macrophage cells. A proportions Z-test and its according p-value are used to provide statistical significance to this finding.

**Figure A2:**
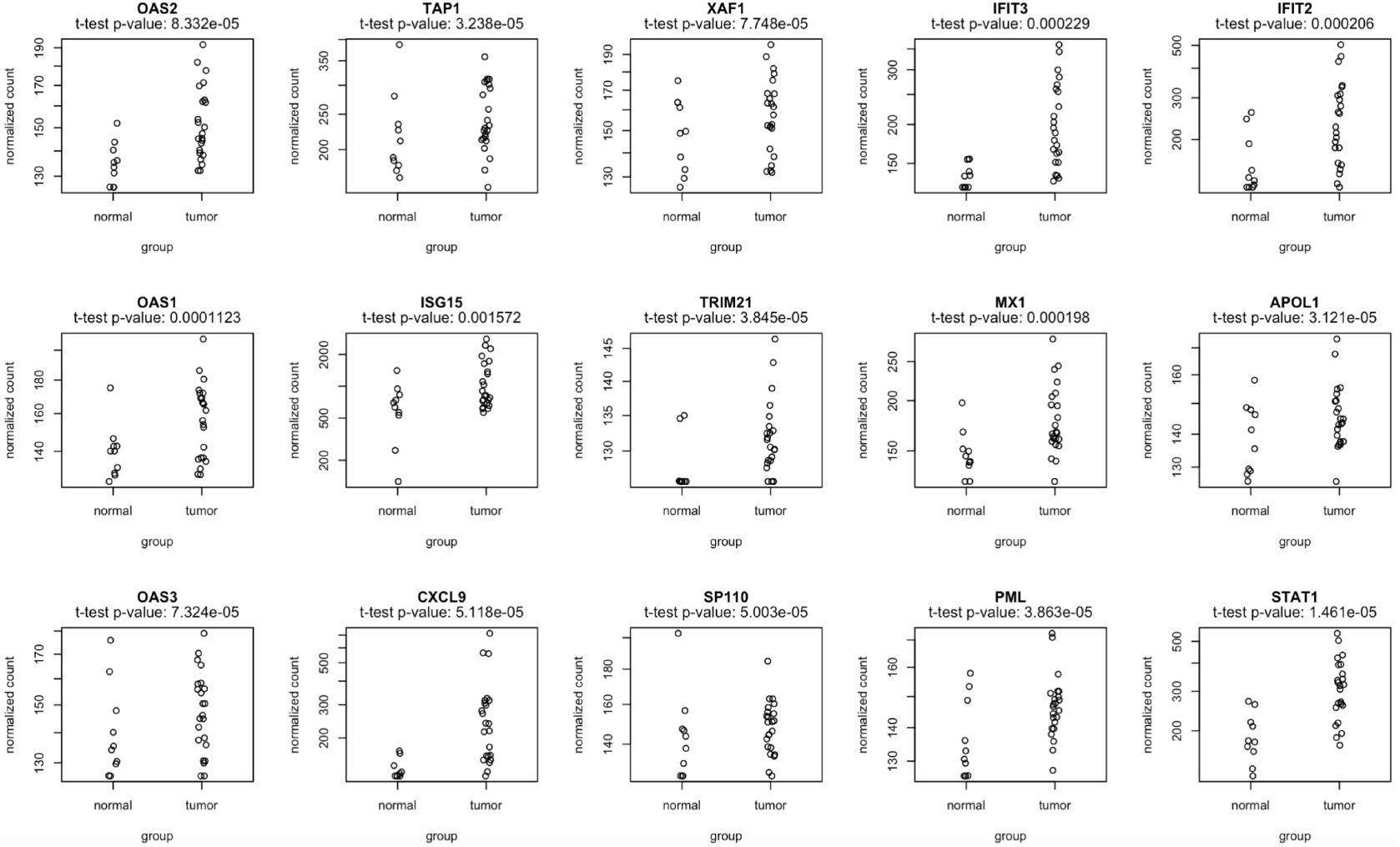
Differential Gene Expression (DGE) analysis for our refined immuno-reactive gene signature. These set of plots showcase the normalized read counts for each gene in our signature in a bulk RNA-seq representation of all cells in our training dataset. We use a t-test p-value to report significance in the up-regulation of genes in the tumor samples.

**Figure A3:**
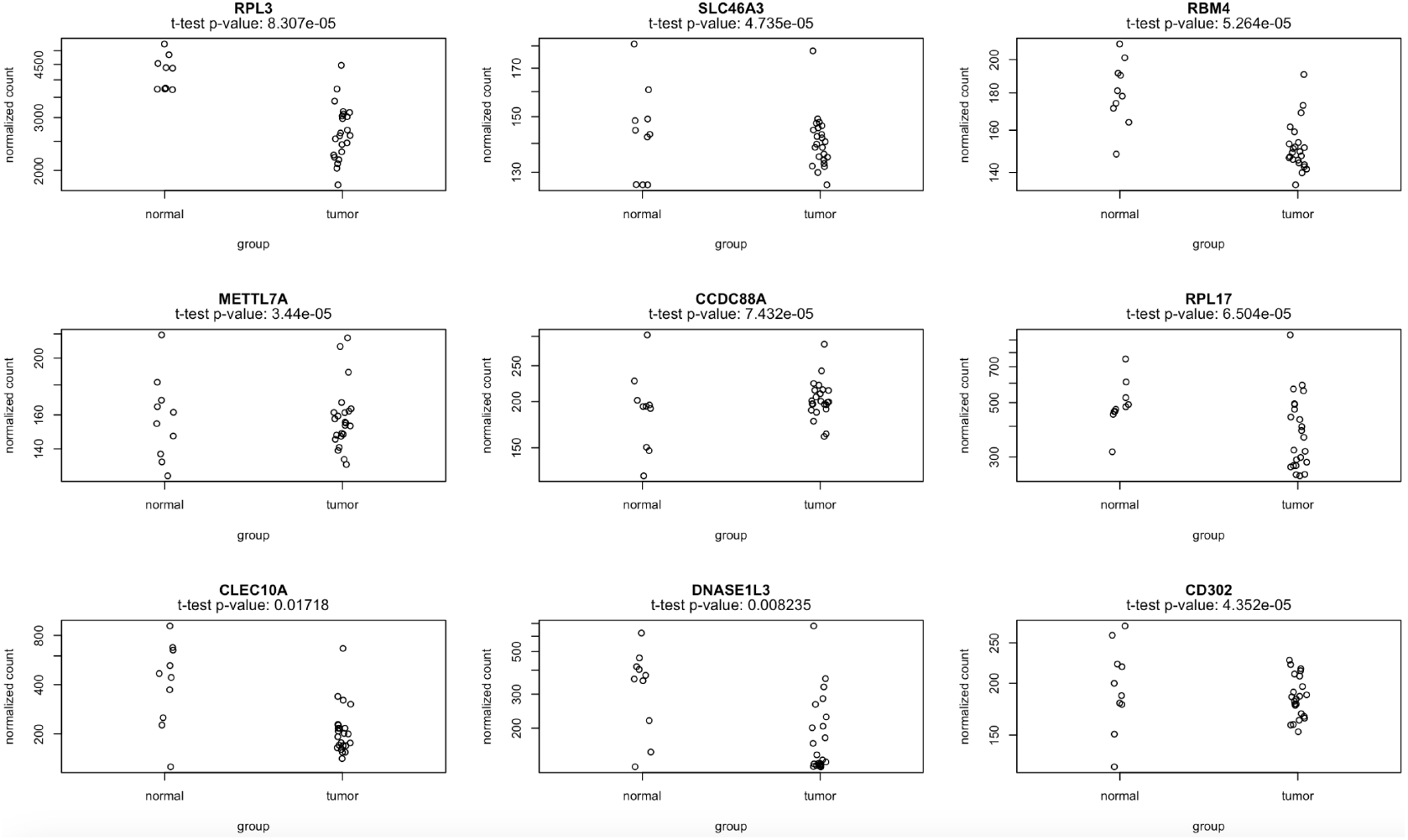
Differential Gene Expression (DGE) analysis for our refined immuno-tolerant gene signature. These set of plots showcase the normalized read counts for each gene in our signature in a bulk RNA-seq representation of all cells in our training dataset. We use a t-test p-value to report significance in the up-regulation of genes in the normal samples.

